# MNC-RED A Chemically-Defined Method to Produce Enucleated Red Blood Cells from Adult Peripheral Blood Mononuclear Cells

**DOI:** 10.1101/616755

**Authors:** Shouping Zhang, Emmanuel N Olivier, Zi Yan, Sandra Suzuka, Karl Roberts, Kai Wang, Eric E Bouhassira

## Abstract

Many methods have been developed to produce red blood cells *in vitro* but translational applications have been hampered by the high cost of production. We have developed R6, a chemically-defined, albumin-free, low-transferrin culture medium, and MNC-RED, a protocol to differentiate peripheral blood mononuclear cells into enucleated erythroid cells that does not require any albumin or any animal components. Erythropoiesis requires large amounts of iron for hemoglobin synthesis. In all existing protocols, these large iron needs are met by increasing the concentration of holo-transferrin. This is necessary because transferrin recycling does not take place in existing erythroid culture conditions. In the R6 medium, iron is provided to the differentiating erythroblasts by small amounts of recombinant transferrin supplemented with FeIII-EDTA, an iron chelator that allows transferrin recycling to take place in cell culture. As a result of the absence of albumin and the use of low amounts of transferrin, the production of cultured red blood cells using the MNC-RED protocol is much less expensive than with existing protocols. The MNC-RED protocol should therefore help make the many translational applications of cultured RBCs economically more feasible.

**Highlights:** We have developed R6, a chemically-defined, albumin-free low-transferrin culture medium, and MNC-RED, a protocol to differentiate peripheral blood mononuclear cells into enucleated erythroid E

R6 is suitable for red blood cell culture despite the low transferrin amounts because of the presence of FeIII-EDTA, an iron chelator that allows transferrin recycling to take place in cell culture.

The MNC-RED protocol should help make the many translational applications of cultured RBCs more economically feasible.

## Introduction

The first liquid culture method to produce red blood cells was developed by Fibach and colleagues who described a two-step procedure in which cytokines were provided in serum and conditioned medium. This method yielded mature erythroblasts but very low rates of enucleation (Fibach et al., 1989).

Since this early work, many authors have shown that hematopoietic progenitors could be dramatically expanded in liquid culture into mature erythroid cells using a variety of recombinant cytokines and small molecules including SCF, EPO, IL3 FLT-3L, IGF1, hydroxy-cortisone and dexamethasone (Dex). Work by von Lindern et al. (1999) and Panzenböck et al. (1998a) revealed that erythroid progenitors present in adult or cord peripheral blood have a limited self-renewal potential since they can be induced to proliferate for about 15 and 30 days, respectively, when cultured in the SED cocktail composed of SCF, EPO and Dex. These same groups also demonstrated that some of these erythroblasts could terminally differentiate into enucleated RBCs when the SCF, EPO and Dex were removed. More recently, the Palis lab demonstrated that the SED cocktail could induce proliferation of primary erythroid precursors isolated from early mouse embryos erythroblasts for more than 80 days (England et al., 2011) and the Bouhassira lab showed that erythroblasts derived from embryonic stem cells could proliferate for more than 60 days (Olivier et al., 2012). Together, these data demonstrate that the SED cocktail is a powerful tool to expand erythroid cells in culture. They also demonstrate that the proliferation potential of erythroid progenitors varies during development, since erythroblasts derived from human embryonic cells are able to proliferate for at least 60-80 days, while those derived from adult peripheral blood expand for less than three weeks.

Using a variation of the SED cocktail, the Migliaccio lab has developed a procedure termed HEMA that yields high numbers of human cultured Red Blood Cells (cRBCs) that are antigenically very similar to RBCs produced in vivo (Migliaccio et al., 2010).

The Douay group demonstrated that very high levels of enucleation could be obtained using a three stage liquid culture system based on recombinant cytokines if the final stages of differentiation were performed on a feeder layer such as mouse MS-5 and human mesenchymal stem cells (hMSC) (Giarratana et al., 2005). This was an important milestone but co-culture on a feeder layer is difficult to scale up. Subsequently, Miharada and colleagues reported that high rates of enucleation of cord blood-derived cRBCs could be obtained without the use of feeder layers by optimization of the culture conditions (Miharada et al., 2006). Building on this work, Timmins and colleagues, recently obtained 90% enucleation and very high yields corresponding to 500 units of cRBCs per initial unit of cord blood in a stirred bioreactor setting using a simple dilution feeding protocol and a humanized medium containing SCF, EPO and IL3 (Timmins et al., 2011). Together, these studies provided a proof of principle that producing RBCs *in vitro* could be economically viable.

The focus of the field therefore shifted to finding a source of immortal cells to produce an unlimited number of red blood cells while cutting costs as much as possible to allow for large scale production (Bayley et al., 2018; Rousseau et al., 2014).

Albumin and transferrin (Tf) are among the most expensive components in erythroid culture media (Rousseau et al., 2014). Albumin is generally purified from either human or bovine serum and is required as carrier and as a source of small molecules. Eliminating albumin from the medium could decrease cost and simplify scaling up because not all batches of albumin support enucleation for reasons that are not entirely clear.

Erythroid cells require large amounts of iron to synthesize heme, the oxygen-binding component of hemoglobin (Byrnes et al., 2016). In most erythroid differentiation protocols, iron needs have been addressed by increasing the concentration of holo-Tf to between 200 and 500µg/mL. This is effective but not particularly efficient because Tf is expensive and is not currently recycled in cell culture conditions.

To decrease costs, culture methods must be simple and reproducible, which is best achieved by using chemically defined components that remove the variation in the quality of animal or human derived reagents that has been a recurring problem for us and many other labs (Ieyasu et al., 2017; Ng et al., 2008; Pei et al., 2017; Uenishi et al., 2014).

We report here that we have developed a new cell culture medium termed R6, which contains no albumin, a low amount of recombinant Tf, as well as MNC-RED, an albumin-free, low Tf, chemically-defined protocol to differentiate human peripheral blood mononuclear cells into large numbers of enucleated RBCs.

## Results

In our standard protocol for terminal erythroid differentiation (condition 1 in Figure 1), human peripheral blood mononuclear cells (MNCs) are expanded for two weeks in albumin-containing StemSpan medium in the presence of hydroxy-cortisone, Stem Cell Factor (SCF), Erythropoietin (EPO) and a combination of Fms-Like Tyrosine kinase 3 Ligand (FLT3L), Interleukin 3 (IL3) and Insulin-like Growth Factor 1 (IGF1) (see week1 and week2 cytokine cocktails in Table S1), yielding day-14 erythroblasts, which express CD71 and CD36 at more than 95% and CD235a at about 50%, and that are morphologically mostly at the pro- and basophilic stages of differentiation (Figure S1).

**Figure 1:**
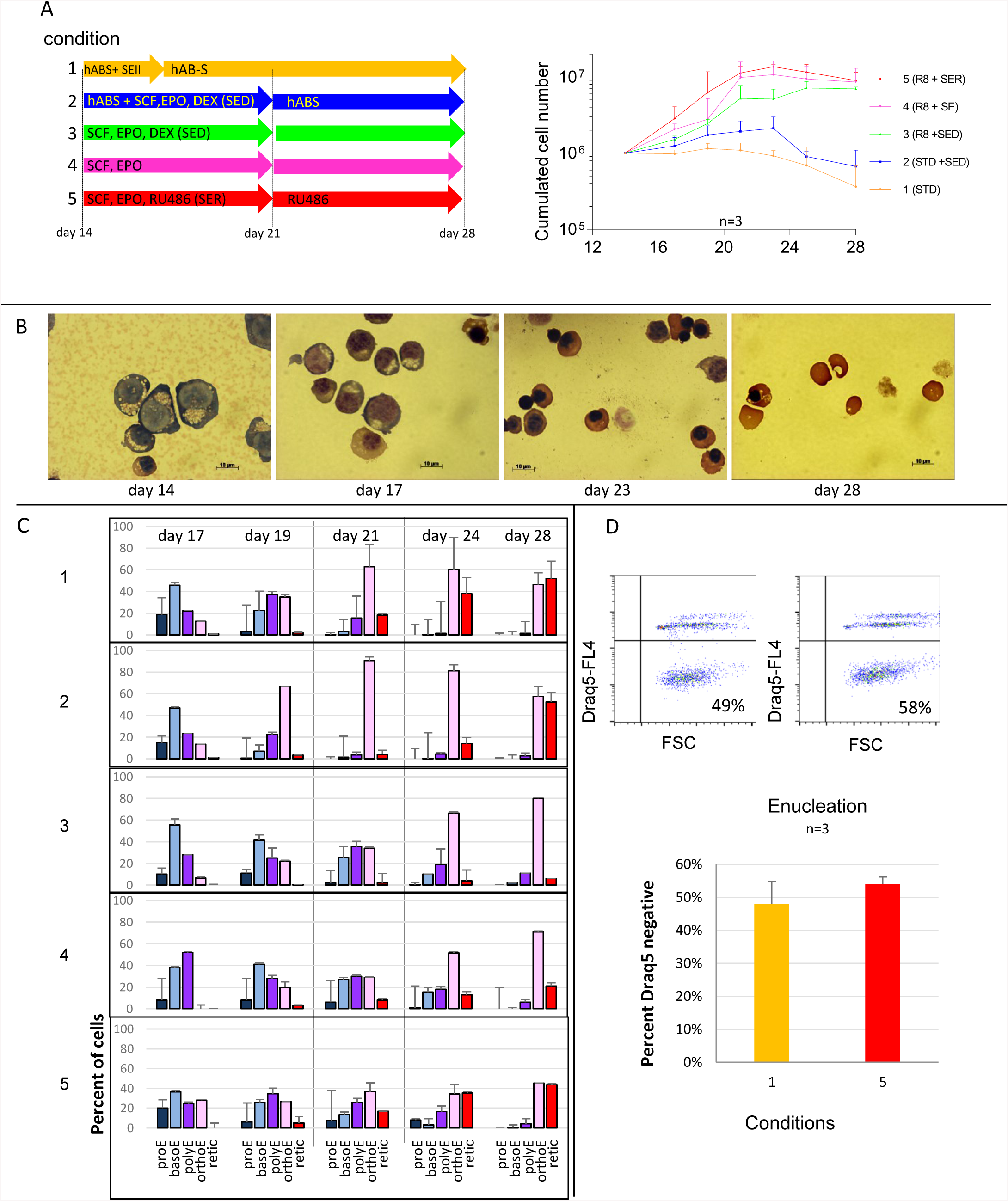
Development of R6 medium. **A:** 2.10^6^ day-14 erythroblasts, that had been obtained by growing PB MNCs for 7 days in week1 medium followed by 7 days in week2 medium, were plated for 14 days either in our standard human AB-serum-containing protocol (condition 1), or in 4 other conditions (conditions 2 to 5) as depicted in diagram on the **top left**. Details of the culture conditions are provided in the method section. Composition of the cytokine and small molecule cocktails (SEII cytokine, SED, SE, SER and R) is provided in Table S1. The line graph on the right summarizes the average and standard deviation (n=3) of the number of live cells observed during the culture as determined by using an hemocytometer after trypan blue staining. Day-14 erythroblasts grown in condition 5 proliferated 10- to 30-fold more than in our standard conditions (condition 1) **B: Micrograph illustrating the morphology of the cells at different times during the differentiation** Micrograph illustrating the typical morphology of the erythroid cells observed at day 14, 17, 23 and 28 of culture after rapid Romanovsky staining. The majority of cells were pro- or basophilic erythroblasts at day 14, basophilic or polychromatophilic erythroblasts at day 17, orthochromatic erythroblasts at day 23 and orthochromatic erythroblasts and reticulocytes (young enucleated RBCs) at day 28. **C: Graph illustrating the quantification of the erythroid differentiation in the conditions depicted above.** About 100,000 cells were cytospined on microscope slides, stained with the rapid Romanovsky method and the cells were classified according to their morphology. proE = Pro-Erythroblasts; basoE = Basophilic Erythroblast.; polyE = Polychromatophilic Erythroblast; orthoE = Orthochromatophilic Erythroblast; retic = reticulocytes (enucleated cRBCs). Graphs illustrate the frequency (average ± SD) of each erythroid precursor during the course of the experiments (n=3). Differentiation in condition 3 was significantly retarded relative to differentiation in condition 2, as illustrated by the lower amounts of orthochromatic and retic and the higher amount of basoE and polyE at day 21 in condition 3. P- and q-values are provided in Figure S2B. Enucleation in condition 3 was very poor (retic < 5%) but was partly restored (21% retic) in condition 4 and completely restored (44% retic) in condition 5. These differences were statistically significant (Figure S2B). **D: Enucleation as measured by flow cytometry:** To confirm the rate of enucleations, the percent enucleation at day 28 was also determined by flow cytometry using Draq5 staining on cells that had been grown in conditions 1 and 5. Rates of enucleation determined by this method were not significantly different from the rates measured after Romanovsky staining.

The day-14 erythroblasts are then seeded in RIT (RPMI, insulin, Tf, ethanolamine, lipids, beta-mercapto-ethanol, lipids) in the presence of 10% human AB-serum and SEII cytokine cocktail (EPO, and low amounts of SCF, IGF-1 and IL3, see SEII cytokines cocktail (Table S1)) for three days, and subsequently incubated for 10-15 days in the same media and serum but without added cytokines. In these conditions, the day-14 erythroblasts enucleate at a rate of about 50-80% (condition1, Figures 1A-C).

This standard protocol has two major issues. Firstly, the number of enucleated cRBCs produced is 50% or less of the number of input 14-day erythroblasts because of massive cell loss after day 21. Secondly, human serum or albumin is required which is problematic because these components are expensive and undefined.

As discussed in the introduction, the maximal period of time during which human peripheral blood erythroid progenitors can be expanded in the presence of glucocorticosteroids, SCF and EPO is two to three weeks (Migliaccio et al., 2002; Panzenböck et al., 1998b). To determine if our day-14 erythroblasts had reached their maximal glucocorticosteroid-induced proliferation, we attempted to amplify them further using the SED cocktail. We first added the SED cocktail to the RIT medium supplemented with human AB serum and cultured the day-14 erythroblasts for 7 days. Terminal differentiation was then induced as in condition 1 by culturing the cells for 7 additional days after withdrawal of the SED cocktail. In these conditions (condition 2) there was a small increase in cell yield at day 21 that did not reach statistical significance. In addition, the number of live cells decreased sharply during days 21 to 28 after withdrawal of the SED cocktail. As a result, the overall yield of enucleated cRBCs was not significantly improved as compared to our standard protocol (Figure 1A). Examination of cell differentiation after rapid Romanowsky staining revealed no significant difference between cells grown in conditions 1 or 2 (Figures 1B and 1C), suggesting that incubation of the day-14 erythroblasts in SED for 7 days did not significantly affect their differentiation when human AB-serum was present.

We then incubated day-14 erythroblasts in SED but changed the base medium from RIT + AB-serum to R8, a serum- and albumin-free medium based on RPMI that we modeled on the E8 media originally developed to grow pluripotent stem cells (Chen et al., 2011). In these conditions (condition 3), the cell yield was significantly different from that of the standard conditions at every time point that we tested (Figures 1A and S2A). The yield of cells increased about 8-fold at day 21 and there was no major decrease in cell number between day 21 and 28. Examination of differentiation after Romanowsky staining revealed that incubation in the SED cocktail in the absence of serum retarded the differentiation since there were significantly more pro- and basophilic erythroblasts and less polychromatic and orthochromatic erythroblasts between day 21 and 28 (Figures 1C and S2B). In addition, enucleation was completely blocked with orthochromatic erythroblasts accumulating at day 28 and eventually dying by day 32-35.

In summary, these experiments revealed that day-14 erythroblasts retained the capacity to divide a few times without differentiating when stimulated with SED without serum. They also revealed that enucleation was very ineffective with or without serum because of the large decrease in cell numbers observed after removal of the cytokine in the former case, and because of the enucleation block in the latter case.

To determine if one of the components of SED might cause the block in enucleation, we set up a series of experiments in which we systematically omitted one or two components of the SED cocktails. This revealed that day-14 erythroblasts grown in R8 containing either EPO, SCF, or Dex alone, EPO and Dex, or SCF and Dex did not grow well and/or died rapidly (Figure S3). However, when the same cells were grown without Dex but with SCF and EPO (condition 4), we observed significant differences with condition 1 since the yield was up to 30-fold higher (Figures 1A and S2A), and enucleation was restored at a rate of about 20% (Figures 1C and S2B).

These experiments suggested that the enucleation block that we observed in condition 3 might be induced by the presence of Dex. Adding RU486 to SCF and EPO (condition 5) confirmed this observation since this glucocorticosteroid antagonist had no major effect on proliferation but restored enucleation levels to that of our standard conditions (Figures 1C, S2A and S2B).

To confirm this result, enucleation was also assessed by flow cytometry, a method that is somewhat more accurate than manual enumeration after Romanovsky staining because many more cells can be evaluated. These experiments confirmed that there were no significant differences in the rate of enucleation between conditions 1 and 5 (Figure 1D).

We concluded from these experiments that the yield of enucleated cRBCs obtained from day-14 erythroblasts can be improved by inducing their terminal differentiation by growing them in EPO and SCF in the absence of Dex for 7 days prior to removing all cytokine stimulation.

To refine our protocol, we titrated some of the components of medium R8 to determine if they were necessary. Experiments in which single component were removed from R8 suggested that ethanolamine at 1.6mM was associated with a small but significant decrease in yield, that lipids and L-ascorbic acid had a small positive effect and that sodium selenite had no effect (Figures S4 and S2C). To confirm these results, we formulated medium R5 which contains only RPMI, lipids, Tf, insulin and L-ascorbic acid and verified the above results by either removing either the L-ascorbic acid, the lipids or by adding sodium selenite. This confirmed that the presence of L-ascorbic acid and lipids was associated with a significantly higher yield, and that sodium selenite had no effect (Figures S4 and S2C). Finally, we directly compared the R5 and R8 media. This revealed that R5 yielded significantly more cells than R8 and that both media yielded similar rates of enucleation (Figures 2 and S2C).

**Figure 2:**
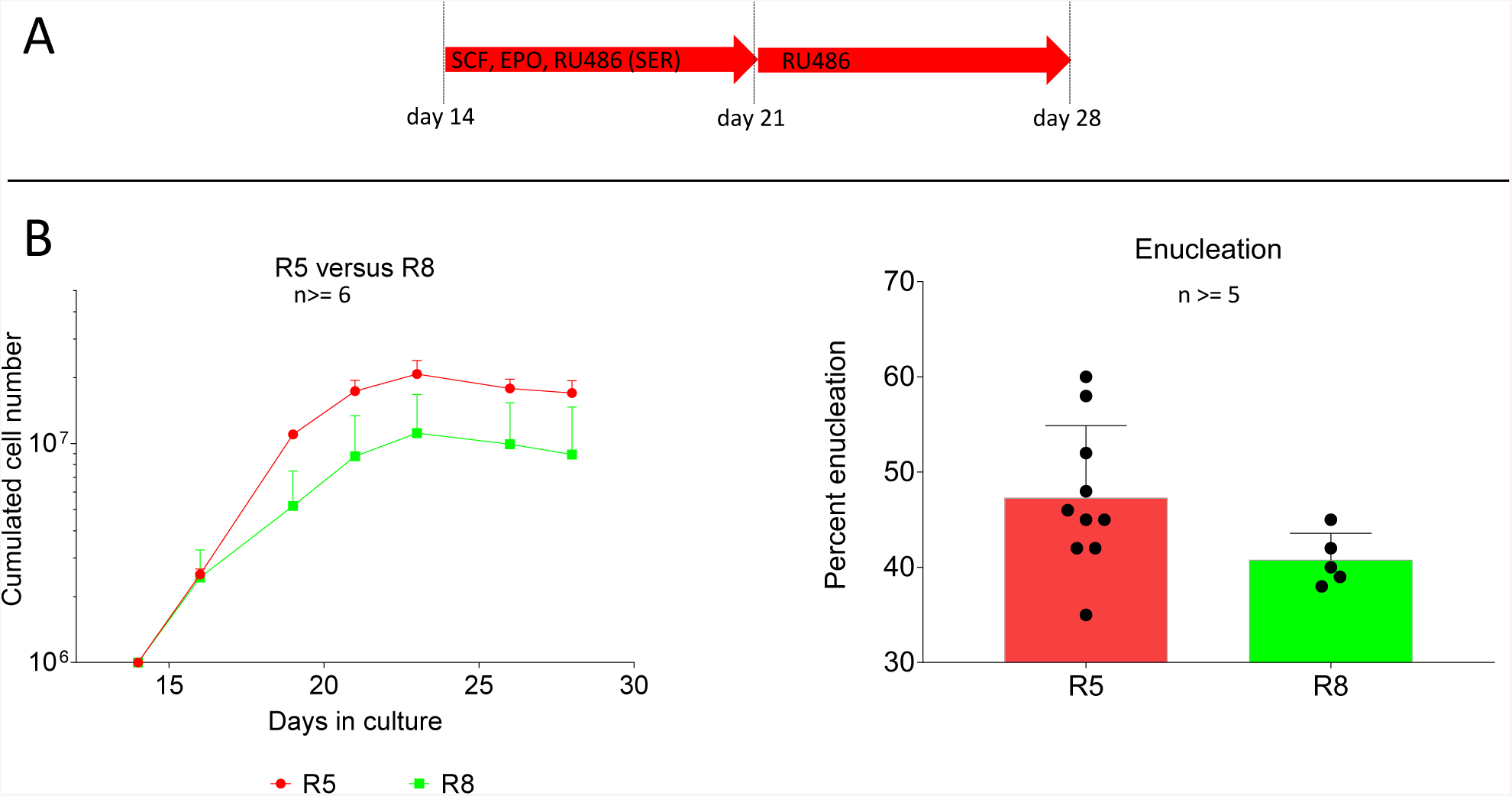
Comparison of R5 and R8 media. **A:** Diagram illustrating the cytokine cocktails used in the experiments. **B:** 3.10^6^ day-14 erythroblasts, were plated for 7 days in either R5 or R8 in the SER cytokine cocktail for 7 days followed by 7 days in R5 or R8 plus RU486. Graph on the left illustrates the number of live cells. The yield of cells was significantly greater in R5 at every time point that were monitored (n= 5). Student t-test p-values - q-values for an FDR of 5% are provided below the graph. Graph on the right illustrates the number of enucleated cells determined after Romanovsky staining. There was no significant difference between the two media.

The experiments described in Figure 1 had shown that RU46 was necessary to help restore enucleation when added at day 14. However, since there is no addition of glucocorticosteroids after day 14, we wondered in RU486 was still necessary after day 21. To test this hypothesis, we compared cell yields and rates of enucleation with or without RU486 added after day 21. This revealed that this compound was not necessary after day 21 (not shown).

### Iron chelators

*In vivo*, holo-Tf is internalized after binding to its receptor, re-exported from the cells as apo-Tf, reloaded with iron, and eventually re-utilized (Luck and Mason, 2012). However, *in vitro*, apo-Tf recycling does not occur in erythroid cell culture for a lack of proper iron source, which is complicated by the poor solubility of iron in most media.

To decrease cost, we tested the hypothesis that providing an appropriate source of iron would enable Tf recycling to occur and therefore allow the culture of erythroid cells in low amounts of Tf.

Titration experiments revealed that the yield of cRBCs when day-14 erythroblasts were grown in R5 was significantly reduced when the concentration of holo-Tf was 50 µg/mL or below, suggesting that at that concentration iron become limiting in these cultures (Figures 3A and S2D).

**Figure 3:**
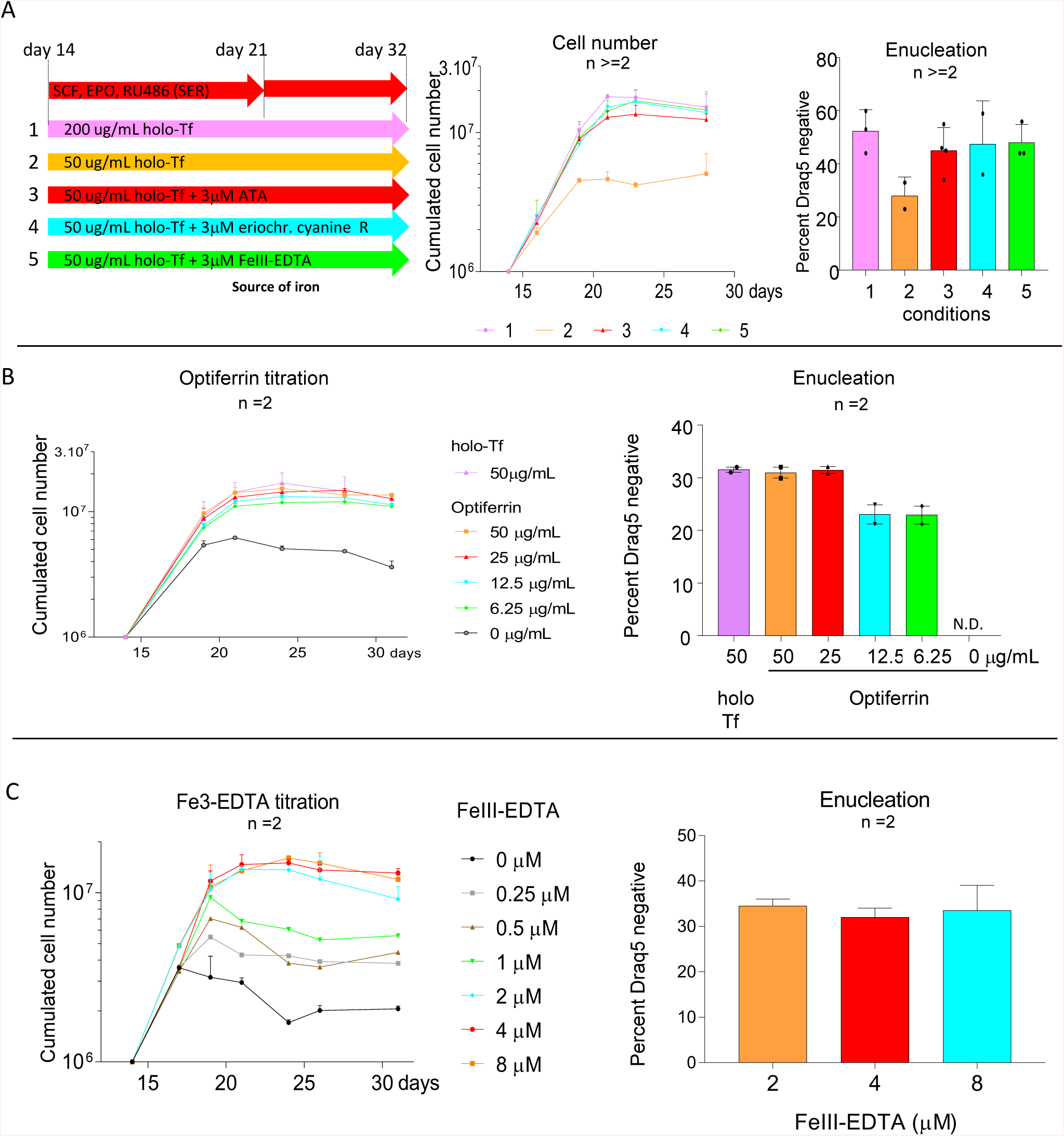
Iron source. **A: Iron chelators.** left: Diagram illustrating the cytokine cocktail used in the experiments (top row) and the source of iron in conditions 1 to 5. Middle: 3.10^6^ day-14 erythroblasts, were plated for 7 days in either R5 medium supplemented with the SER cytokine cocktail for 7 days followed by 7 days in R5 medium alone. The graph illustrates the number of live cells at different days in the culture in the presence of different amounts of human holo-Tf and different iron chelators. Cell numbers were determined using a hemocytometer after trypan blue staining. Cells grew poorly with only 50 µg/mL of holo-Tf in the medium (compare conditions 1 and 2). Cell growth in medium containing 50µg/mL of holo-Tf was completely restored in the presence of all FeIII-EDTA and eriochrome cyanine R and partially restored in the presence of ATA (n=3, p- and q-value in Figure S2D). Right: graph illustrating enucleation (determined by flow cytometry) in the 5 conditions tested. There were no significant differences in the rates of enucleation (n=2). **B. Optifferin titration.** 3.10^6^ day-14 erythroblasts were cultured with the same cytokines as in A in R5 supplemented with 3µM FeIII-EDTA and the indicated concentrations of Optiferrin. Graph on the left illustrates the number of live cells at different days in the culture in the presence of different amounts of Optiferrin. Concentration of Optiferrin as low as 6.25 µg/mL supported the growth of day-14 erythroblasts (n=2, p-and q-value are in Figure S2E). Graph on the right illustrates rate of enucleation in the different conditions tested. Rate of enucleation were not significantly different (n=2). We decided to use 20 µg/mL of Optiferrin in subsequent experiments **C: FeIII-EDTA titration.** 3.10^6^ day-14 erythroblasts were cultured with the same cytokines as in A in R5 supplemented with 20 µg/mL of Optiferrin and the indicated concentration of FeIII-EDTA. Graph on the left illustrates the number of live cells at different days in the culture in the presence of different amounts of FeIII-EDTA. The graph on the right illustrates the rates of enucleation. There were no significant differences between the yield of cells nor the rate of enucleation in the presence of 2, 4 or 8 µM FeII-EDTA (n=2). Cells grew poorly in the presence of 1 µM or less of FeIII-EDTA. We decided to use 4 µM FeIII-EDTA in subsequent experiments.

To find an appropriate source of iron, we compared the yield of cells and the rate of enucleation when day-14 erythroblasts were grown in R5 containing either 200µg/mL of holo-Tf, or 50µg/mL of holo-Tf in the presence of a mixture of 220nM Fe(NO_3_)_3_ and 3.2µM Fe_2_(SO_4_), the classical source of iron in erythroid culture (Giarratana et al., 2005), or in the presence of iron chelators such as iron citrate, aurintricarboxylic acid (Bertheussen, 1993) (ATA), eriochrome cyanine R (Kuban-Jankowska et al., 2016) or FeIII-EDTA (Aasa et al., 1963).

This revealed that the ferric nitrate/ferrous sulfate mixture and the iron citrate were not effective as a source of iron (data not shown). By contrast, the three other iron chelators were effective. Supplementation of 50µg/ml Tf with either 3 µM eryochrome cyanide or FeIII-EDTA completely restored cell growth since we observed no significant difference with the control culture containing 200µg/mL of holo-Tf (Figures 3A and S2D). In the case of 3µM ATA, the rescue was only partial, since the yield was higher than with no supplementation, but significant different from that of the 200µg/mL holo-Tf control (Figures 3A and S2D). Additional experiments suggested that combinations of the three chelators had no advantage over the use of a single chelator (data not shown). Although there was no significant difference between eriochrome cyanine R and FeIII-EDTA, the latter compound had yielded the highest average number of cells. We consequently decided to focus on that molecule for the rest of the study.

### Optiferrin

Holo-Tf is purified from human plasma. It is therefore a potential source of variation. To determine if it could be replaced by Optiferrin, a commercially available recombinant Tf produced in rice, we cultured day-14 erythroblasts in the presence of 3µM FeIII-EDTA and either 50 µg/mL holo-Tf or decreasing amounts of Optifferin. This revealed that there was no significant difference between the yield of cells obtained with 50 µg/mL of holo-Tf and as little as 12.5 µg/mL of Optiferrin (Figure 3B). The day-14 could also grow in 6.25 µg/mL of Optiferrin but the yield was significantly lower at day 28 (Figures 3B and S2E). Enucleation rates in the presence of 50 µg/mL of holo-Tf or at least 12.5 µg/mL of Optiferrin were not significantly different. To maintain an excess of Optiferrin at all times in these cultures, we decided to use a concentration of 20µg/mL in subsequent experiments.

To determine the optimal concentration of FeIII-EDTA, we titrated it between 0µM and 8 µM. This revealed that the yield of cRBCs and the enucleation rates were not significantly different when the day-14 erythroblasts were cultured in the presence of 2, 4 or 8µM of FeIII-EDTA (Figure 3C). By contrast the day-14 erythroblasts grew poorly when the concentration of FeIII-EDTA was below 2µM. An additional experiment revealed that there was no obvious toxicity when the concentration of FeIII-EDTA was increased to 20µM (not shown). We decided to use 4µM FeIII-EDTA in future experiments. We therefore reformulated medium R5 by replacing the 200µg/mL holo-Tf with 20µg/mL Optiferrin and 4µM FeIII-EDTA, creating medium R6 which is significantly cheaper to produce than R5 or R8.

### Tf recycling

To determine if, as hypothesized, the positive effect FeIII-EDTA was associated with Tf recycling, we directly measured Optiferrin iron loading by Poly-Acrylamide Gel Electrophoresis (PAGE) in 6M urea as described previously (Makey and Seal, 1976). We first demonstrated that FeIII-EDTA could deliver iron to Optiferrin by preparing R5 medium containing 20µg/ml of Optiferrin, as provided by the manufacturer and either no additional iron, 0.22µM Fe(NO_3_)_3_ and 3.2µM Fe_2_(SO_4_), or various amounts of FeIII-EDTA. After 2 hours of incubation at 37C the media were concentrated and 2µg of Optiferrin was examined by PAGE (Figure 4). Staining with Coomassie blue revealed that the Optiferrin provided by the manufacturer was almost completely in the apo conformation, but that it was fully loaded with iron in the presence of 4 or 20 µM of FeIII-EDTA, and partially loaded with 2µM of the iron chelator (Figure 4). The combination of Fe(NO_3_)_3_ and Fe_2_(SO_4_) was completely ineffective to load Optiferrin with iron in the context of the R6 medium explaining why these compounds are not effective as a source of iron in erythroid culture.

**Figure 4.**
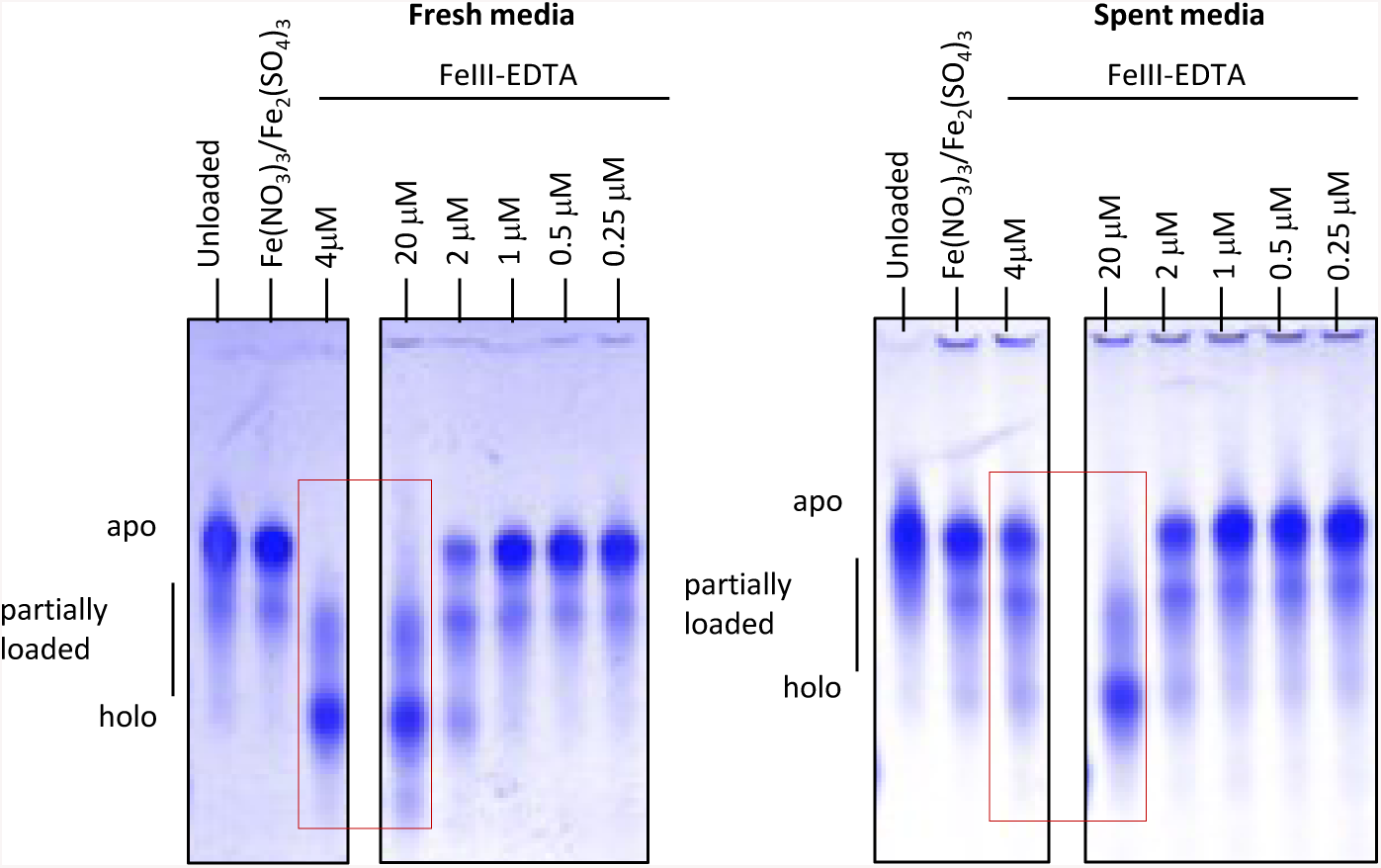
Optiferrin recycling. Fresh or spent R6 media containing 20µg/mL of Optiferrin and various amounts of FeIII-EDTA was analyzed on a 6M urea 5% poly-acrylamide gel. In fresh media (left panel), Optiferrin was in the apo form in the absence of iron, in the presence of Fe(NO_3_)_3_/Fe_2_(SO_4_) (0.22µM/3.2µM), or with less than 2µM FeIII-EDTA, but was in the holo form in the presence of 4 or 20µM FeIII-EDTA, demonstrating that FeIII-EDTA can deliver iron to Optiferrin. In the spent media (right panel), Optiferrin was in the apo form in the presence of 4µM of FeIII-EDTA suggesting that the iron was depleted, but was still in its holo-form in the 20µM condition, suggesting that iron recycling can occur in the R6 media. Gels are representatives of the results of two independent experiments.

To determine if Optiferrin could be recycled in the presence of FeIII-EDTA, about 2 million day-14 erythroblasts were incubated for four days without any feeding in R6 media containing various amounts of iron (and 20µg/mL of Optiferrin). As expected, cells proliferated in all conditions containing at least 2µM FeIII-EDTA. The spent media was then recovered, concentrated and analyzed by 6M urea 5% polyacrylamide gel electrophoresis as above. This revealed that Optiferrin in media containing 4µM or less of FeIII-EDTA was now almost completely in its apo form, suggesting that the iron had been completely consumed by the cells (Figure 4). By contrast, Optiferrin in the medium containing 20 µM of FeIII-EDTA was still in its holo form demonstrating that FeIII-EDTA can be used as a source of iron to allow Optiferrin recycling to take place in erythroid cell cultures.

### MNC-RED protocol

All of the above experiments were performed with day-14 erythroblasts that had been generated by expansion in albumin-containing StemSpan medium of MNCs derived from a single individual. To determine whether the R6 medium was effective on MNCs from other individuals, and whether MNCs could be differentiated into cRBCs using a fully albumin-free, chemically-defined protocol, we designed the MNC Robust Erythroid Differentiation protocol (MNC-RED, Figure 5A). In this protocol MNCs are differentiated into day-14 erythroblasts using the same cytokine cocktails as in our standard protocol, but StemSpan is replaced with IMIT, a chemically-defined medium that we recently developed (see accompanying manuscript). IMIT contains methyl-β-Cyclodextrin, Trolox, Insulin, Optiferrin, FeIII-EDTA, Lipids and Ethanolamine (Table S1). The day-14 erythroblasts are then differentiated into enucleated RBCs in R6 medium using the optimal conditions described above (see methods).

**Figure 5:**
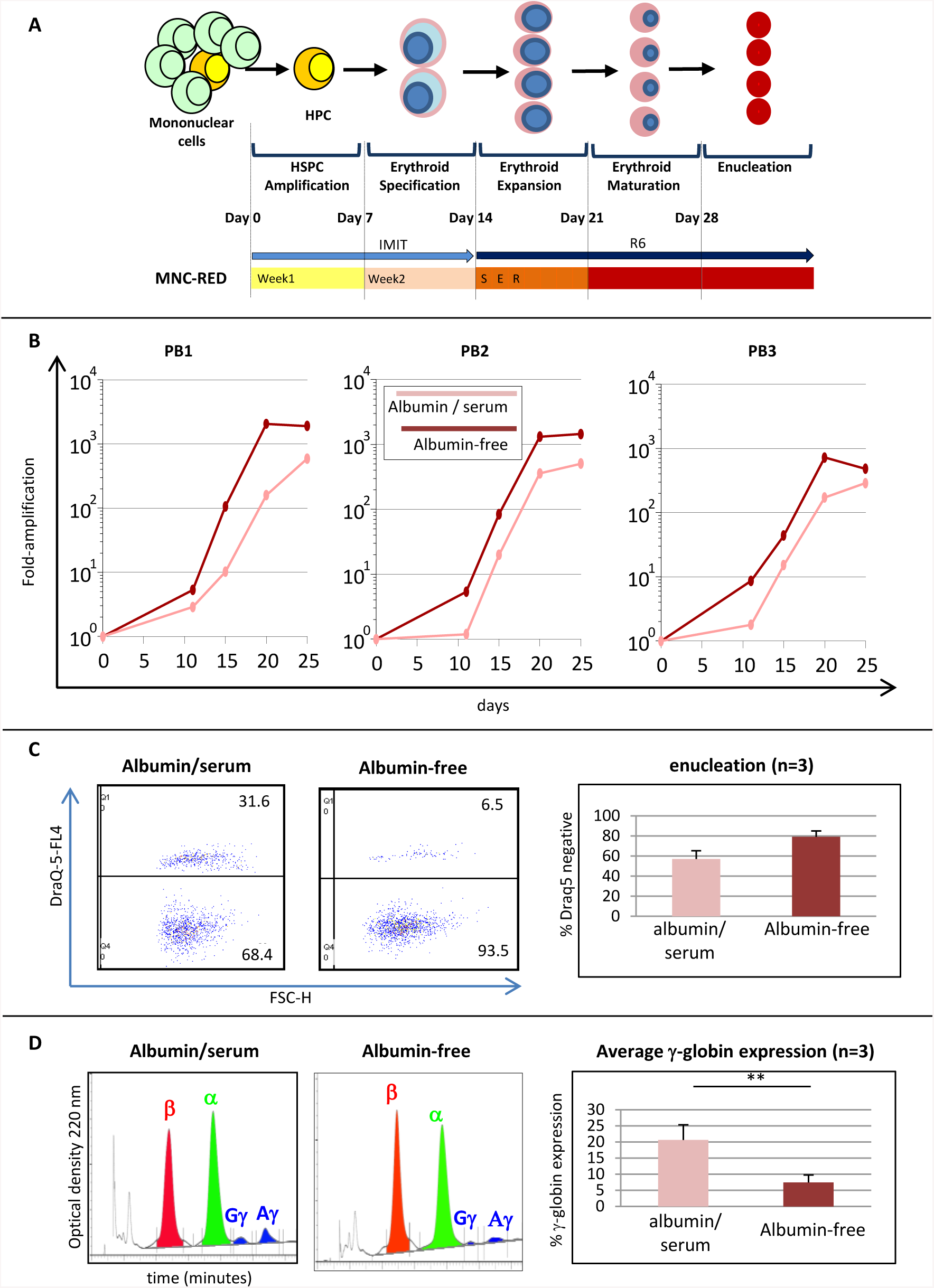
MNC-RED protocol to generate enucleated cRBCs from PB mononuclear cells. **A:** Graph summarizing the MNC –RED protocol. Media and supplement composition can be found in Table S1. **B**: Scatter plots illustrating the growth of PB MNC cells collected from three healthy individuals and incubated for 28 days using the MNC-RED protocol or in albumin-containing conditions using the standard protocol. The yield of enucleated erythroid cells in the MNC-RED protocol was higher for all three individuals. When the data from the three individuals was averaged, the difference between the two protocols was significant (Figure S5) **C**: cRBCs produced using either the albumin-containing or albumin-free protocol were analyzed by flow cytometry on day 32 of the protocol to assess the rate of enucleation using nuclear stain Draq5 (and propidium iodide to exclude the dead cells). FACS plots illustrating the enucleation rate for individual PB1 (measured at day 32). Right: histogram illustrates the rates of enucleation (± SEM) in albumin-free and albumin-containing conditions averaged for the three individuals (n=3). For all three individuals, cells grown using the MNC-RED protocol exhibited a higher rate of enucleation but the difference did not reach significance. **D**: Chromatograms illustrating globin expression in the cRBCs derived from individual PB1 in albumin-containing and albumin-free condition. Right: histogram illustrates the significant difference in the percentage of expression of gamma globin in albumin-free and albumin containing conditions for the three individuals tested (n=3; paired t-test p-value = 0.0345). Percent γ-globin expression was calculated as 100* (Gγ+Aγ)/(Gγ+Aγ+β).

Expanding and differentiating MNCs from three healthy individuals in albumin-free, chemically-defined conditions proved highly efficient (Figure 5B) since the MNC-RED protocol yielded 1,293 ± 739 cRBCs/MNC (average ± SD) as compared to 465 ± 156.5 cRBCs/MNC when cells from the same individuals were grown using our standard protocol. The difference in yield was statistically significant at days 21 and 25 (Figure S5). At day 32, the rate of enucleation with the R6 medium reached 94% in one individual and averaged 79.3 ± 18.46% (average ± SD), as compared to 57.1 ± 20.75% when albumin-containing media were used (Figure 5C). This difference, however, did not reach statistical significance (t-test p-value = 0.23).

To determine if globin expression was affected when MNCs were differentiated using either the MNC-RED or our standard serum-containing protocol, globin expression was analyzed by HPLC. This revealed that the cRBCs produced using MNC-RED expressed mostly Hb A and an average of 7.3 +/- 3.8% Hb F (average ± SD) while the cells produced using our standard protocol expressed 20.6 +/- 8.0 % Hb F. This difference was significant (t-test p-value = 0.008, Figure 5D). We concluded that the cells produced using MNC-RED express lower amounts of Hb F than cells produced with the standard protocol. The former cells are therefore more similar to cells produced *in vivo*, but are still over-expressing Hb F, since *in vivo* produced cells generally express less than 1% Hb F.

## Discussion

We have developed R6 a chemically-defined cell culture medium suitable for erythroid culture. R6 is inexpensive because it contains no albumin and low amounts of Optiferrin. It is nevertheless suitable to produce cRBCs because the presence of FeIII-EDTA allows the recycling of Optiferrin. In combination with our improved terminal erythroid differentiation which involves withdrawing the glucocorticoid stimulation 7 days prior to withdrawing SCF and EPO, the R6 medium yields about 15-30 fold more enucleated cells than our previous protocol.

We do not have a mechanistic explanation for the improved cell yield that we obtained by withdrawing glucocorticoid stimulation before removing the SCF and EPO stimulation to induce terminal differentiation. Bauer et al. have shown that the glucocorticoid receptor is required for stress erythropoiesis but not steady-state erythropoiesis (Bauer et al., 1999). Stellacci et al. have shown that the glucocorticoid receptor directly interacts with the EPO receptor and that combined Dex and EPO stimulation inhibits some but not all of the effects of EPO, which leads to a delay in pro-erythroblasts maturation and erythroid expansion (Stellacci et al., 2009). Eliminating Dex while maintaining EPO and SCF stimulation, in cells that have already been subject to a long exposure to glucocorticosteroids, might allow stress erythroblasts to slowly re-enter the normal erythroid differentiation pathway and resume their terminal differentiation in a more efficient manner than when all three components of the SED cocktails are removed at the same time.

Tf and albumin are expensive and represented 30 to 40% of the total cost of cRBC production because they are required in large amounts until the end of the culture at a time when the culture volumes are maximal (Rousseau et al., 2014). MNC-RED is therefore significantly less expensive than our previous standard protocol. It is also more highly reproducible because of the elimination of all undefined components. Finally MNC-RED can easily be scaled-up into a fed-batch bioreactor system, because it does not require any feeder cells nor any centrifugation, as media changes can be performed by simple medium addition combined with supplementation with RU486 to eliminate any residual glucocorticosteroids.

The MNC-RED protocol will likely be useful for studies that require the production of adult cRBCs, for instance, to study disease mechanisms or for the production of small batches of cRBCs for therapeutic use. It might also help decipher transduction and enucleation mechanisms in late erythroid differentiation because the complete elimination of all undefined components will provide an ideal platform for mechanistic studies.

## Methods

### Samples

Peripheral blood (PB) cells were obtained from healthy donors under a protocol approved by the Albert Einstein College of Medicine IRB.

Mononuclear cells from whole blood were purified using Histopaque (Sigma-Aldrich) according to the manufacturer instruction, and residual RBCs were lysed by addition of 5 volumes of cold (4°C) RBC lysing buffer (NaHCO3 790mg/L and NH4Cl, 7.7g/L) for 10 minutes. Aliquots containing about three million cells were then frozen and stored in liquid nitrogen.

### Reagents

The suppliers for all reagents are provided in Table S2.

### Culture conditions

All cultures were performed at 37C, in 5 or 10% CO_2_ depending on the presence of stable glutamine in the medium. All experiments were performed in 5% O_2_ in accordance with previously published results (Vlaski et al., 2009),

### Differentiation of peripheral blood mononuclear cells into day-14 erythroblasts and into cRBCs

#### Standard albumin-containing protocol

Frozen aliquots of PB mononuclear cells were seeded in the following 4-step protocol: In the first step (days 0 to 7), cultures were initiated at a density of 500,000 cells/mL with serum-free StemSpan SFEM (Stem Cell Technologies) supplemented with Hydrocortisone (1µM), IL3 (7ng/mL), FLT3L (17ng/mL), SCF (50 ng/mL), and EPO (1.3 U/mL) for 7 days.

In the second step (day 7 to 14), the cells were transferred to StemSpan medium supplemented with hydrocortisone (1µM), IL3 (7 ng/mL), SCF (20 ng/mL), EPO (3.3 U/mL) and IGF-1 (20 ng/mL) for 7 days. Cell density was kept below two million cells per mL at all times by adding fresh medium every 2 or 3 days as needed.

In the third step (day14 to 17), the day-14 erythroblasts thus generated were plated in RIT (RPMI, beta-mercapto-ethanol 0.1mM, Ethanolamine 1.6mM, Lipids (1/200), insulin 10µg/mL, holo-Tf 200 µg/mL) in the presence of 10% hAB-serum, and the SEII cocktail (3U/mL of EPO, and low amounts of SCF (4.4 ng/mL), IGF-1 (4.4 ng/mL) and IL3 (1.5ng/mL)) for three days, and subsequently incubated for 10-15 days in RBIT alone.

For all experiments described in Figures 1 to 3, MNCs that had been frozen at day 11 of culture were expanded until day 14 to test variations of the last 2 weeks of the protocol.

In Figure 1, condition 1, the cells were grown exactly as described in the standard protocol above. In condition 2, the SED cocktail was added at day 14 instead of the SEII cocktail and the cells were cultured for 7 days prior to induction of differentiation as in condition 1.

In condition 3, the cells were grown in R8 medium supplemented with the SED cocktail for 7 days followed by 7 days in R8 medium alone.

In condition 4 and in the experiments described in Figure S3, the cells were grown as in condition 3 except that the various components of the SED cocktail were systematically omitted.

In condition 5, the cells were grown as in condition 4 except that 1µM RU486 was added from day 14 to 28.

In Figure S4 the day-14 erythroblasts were grown as in condition 5 of Figure 1 (1 week in SCF, EPO and RU486 (SER) + one week in RU486) but the base media were either R5, R8, or a modification of these media as indicated.

In Figure 3, the cells were grown in R5 medium plus SER for 7 days followed by 7 days in R5 alone (since we had shown at that point in time that RU486 was not necessary after day 21). The source of iron was varied as indicated. Optiferrin was dissolved in water and used as provided by the manufacturer (without pre-loading it with iron).

In Figures 5 and S5, MNCs from three healthy individuals were differentiated into cRBCs using either the standard protocol described above or the MNC protocol described below:

### MNC-RED protocol

Frozen aliquots of PB mononuclear cells were seeded in the following 4-step protocol:

<u>In the first step</u>, cultures were initiated at a density of 5×10^5^ cells/mL and incubated for 7 days in IMIT supplemented with IL3 (7ng/mL), FLT3L (17ng/mL), SCF (50 ng/mL), EPO (1.3 U/mL) and Hydrocortisone (1µM).

<u>In the second step</u>, the cells were replated at a density of 3-5×10^5^ cells/mL and incubated for 7 days in IMIT supplemented with IGF-1 (20 ng/mL), IL3 (7 ng/mL), SCF (20 ng/mL), EPO (3.3 U/mL) and hydrocortisone (1 µM). Cell density was kept below 2×10^6^ cells/mL at all times by diluting the cells to 3-5×10^5^ cells/mL in fresh medium plus cytokines every 2 or 3 days as needed.

<u>In the third step</u>, the cells were replated at 5×10^5^ cells/mL and incubated for one week in R6 supplemented with SCF (50ng/mL), EPO (3U/mL) and 1µM RU486. Cytokines and medium were renewed as above

<u>In the fourth step</u>, cells were replated at 5×10^5^ cells/mL and allowed to enucleate by incubation for 7-12 days in R6 alone (in the absence of RU 486).

### Analysis and characterization

#### Cell enumeration

Cells were counted with a Luna-FL dual channel Automated Cell Counter (Logos) using acridine orange to visualize the live cells and propidium iodide to exclude the dead cells. Alternatively, cells were counted on a hemocytometer after trypan blue staining.

#### Flow cytometry

MNCs undergoing differentiation were evaluated by FACS using antibodies against CD36, CD71 and CD235a, also known as glycophorin A (BD Biosciences and eBioscience).

#### Enucleation

The enucleation rate was measured using the DRAQ5 DNA nuclear stain (ThermoFisher) after exclusion of dead cells with Propidium Iodide. The cells were analyzed with a BD FACS Calibur flow cytometer (BD Biosciences) or a DPX10 (Cytek) flow cytometer and the flow cytometry data were analyzed with the Flowjo software. At least 10,000 cells were analyzed. Alternatively, enucleation was measured by manual enumeration of at least 200 cells after Rapid Romanovsky staining of cytospin preparations.

#### Cytological staining

Erythroid differentiation and enucleation were also assessed microscopically by Rapid Romanovsky staining of cytospin preparations using the HEMA-3 kit from Fisher Scientific as recommended by the manufacturer. Cell sizes were estimated on a Nikon TE-2000S microscope using software provided by the manufacturer. Alternatively, cells were stained with Wright-Giemsa stain from Thermo-Fisher.

#### RBC filtration

To eliminate the nuclei at the end of the experiments, RBCs were filtered using PAL Acrodisc 25mm WBC filters as recommended by the manufacturer. Filtered cRBCs were stored for up to one month with little signs of hemolysis in Alsever’s solution (Sigma).

#### HPLC analysis

Cells were washed twice with PBS, and lysed in water by 3 rapid freeze-thaw cycles in dry-ice and a 37C water bath. Debris was eliminated by centrifugation at 16,000g and the lysates stored at −80°C. HPLC was performed as described (Fabry et al., 2003). Briefly, a few µL of lysate containing about 50µg of protein in about 100 µL of 40% acetonitrile and 0.18% TFA was filtered and loaded on a VYDAC C4 column. The globins were then eluted with increasing concentration of acetonitrile during a period of about 80 minutes. The starting elution buffer was programmed to be 80% buffer A and 20% buffer B and to rise to 50% buffer B in 50 minutes. Buffer A = 36% acetonitrile and 0.18% TFA and buffer B = 56% acetonitrile and 0.18% TFA. Globin chain elution was monitored by measuring O.D. at 220 nm.

#### Transferrin analysis

Tf and Optiferrin iron loading was determined in 6M PAGE as described previously (Makey and Seal, 1976). About 2-5 µg of protein were loaded in each well. When necessary the Tf and Optiferrin were concentrated from the culture medium by centrifugation through 0.5mL 10KD MWCO Amicon ultra-centrifugal filters according to the manufacturer instructions (Merck Millipore).

#### Statistical analysis

Significance was assessed using GraphPad Prism 7 for Windows by performing student t-tests analyzing each row individually. Q-value were calculated using the False Discovery approach (two-stage step-up method of Benjamini, Krieger and Yeketieli as recommended by GraphPad) using a threshold of 5% FDR.

## Acknowledgments

SZ, ENO, ZY, SZ, KR, KW and EEB were supported in part by grants NYSTEM C030135, C029154; NIH HL130764; and Doris Duke Foundation 2017087. We thank Connie Westhoff from the New York Blood Center for providing samples.

## Author contributions

SZ, ZY, SZ, KR, KW performed some of the experiments and contributed to experimental design and data interpretation. ENO performed some the experiments, contributed to experimental design,s ata interpretation and manuscript writing EEB supervised project, contributed to experimental design, data analysis and manuscript writing.

