## Supplementary Figures and Tables Legends for "MNC-RED A Chemically-Defined Method to Produce Enucleated Red Blood Cells from Adult Peripheral Blood Mononuclear Cells"

**Supplementary Figure Legends:**

**Figure S1: day-14 erythroblasts.**

A: Micrograph illustrating the typical morphology of day-14 erythroblasts after rapid Romanowsky staining. Most cells are pro-and basophilic erythroblasts but up to 15% of the cells are more mature polychromatophilic and orthochromatophilic erythroblasts.

B: Flow cytometry. Dotplot illustrating a representative example of expression of CD71, CD36 and CD235a in day-14 erythroblasts. About 98.9% of the cells express CD71, 95.2% CD36 and 58.1% CD235a (n = 2). This data supplement the Romanowsky staining and confirms that most of the cells are committed to the erythroid pathway at this point of time in the culture.

**Figure S2: p-and q-value for Figure 1A, 1C, 2, 3A, 3B and S4.**

t-test p-values were calculated for each pair of conditions. The q-values were calculated using the Two-stage step-up method of Benjamini, Krieger and Yekutieli using a desired FDR of 5%.

P- and q-values were calculated systematically for all combination of conditions. For each experiments, conditions in which none of the row were significant are not shown.

A: In figure 1A, the growth rate in condition 1 was significantly different from that of conditions 3 and 5 at most of the days tested. Condition 2 was significantly different from condition 5 at days 21 to 25; (n = 3)

B: In figure 1C, the proportion of the various erythroid precursor during the time course of the experiment was significantly different at multiple days between conditions 1 and 3, 1and 4, 1 and 5, 2 and 3, 2 and 4 and 2 and 5. The number of enucleated cells (retic) was also significantly different between conditions 3 and 5.

C: In figures 2 and S4, Ethanolamine proved to be toxic (n = 2) while acid ascorbic (n = 5) and lipids (n = 2) improved cell yields. R5 yielded more cells that R8 (n >= 6)

D: **:** In figure 3A, the growth rate of the cells was significantly different at multiple days between conditions (1 and 2), (1 and 3), (2 and 3), (2 and 4) and (2 and 5); (n = 3). We concluded that eryochrome cyanide and FeIII-EDTA could both be used to provide iron in erythroid culture and allow the growth and differentiation of the cells with lower amounts of transferrin.(n = 2)

E: In figure 3B, cells could not grow in the absence of Optiferrin and the yield of cells was lower when the concentration of Optiferrin was 6.25µg/mL. (n = 2)

**Figure S3: Effects of the SED components**

A: Culture conditions: 14-days erythroblasts were grown in R8 for 7 days in the presence of the indicated cytokines followed by 7 days in R8 alone.

B: Growth rates observed in the presence of the SED cocktail, of SCF alone (S), EPO alone (E), Dex alone (D), SCF and Dex (SD) or EPO and Dex (ED). 14-days erythroblasts expanded in the presence of the SED cocktail (see Figure 1A), but very little or not at all, in all other conditions tested in this figure. We concluded that in the absence of serum, 14-day erythroblasts can significantly expand and terminally differentiated without significant loss of cells in the presence of SCF and EPO, but not if either of these cytokines is absent (see main text).

**Figure S4: Analysis of the component of the R8 medium**

Diagram on the top right illustrating the cytokine cocktails used in the experiments. $3.10^6$ day-14 erythroblasts, were plated for 7 days in various base media (derived either from R5 or R8) in the SER cytokine cocktail for 7 days, followed by 7 days in the same conditions but without SCF or EPO. All other graphs illustrate the cumulated number of live cells as a function of days in cultures.

Top left: Cells grown in R8 with 1.6mM ethanolamine grew less rapidly that with no ethanolamine (n=2, q-value <0.001 9FDR < 5%) for days 21 to 28).

Second row: Cells were grown in R8 containing decreasing amount of L-ascorbic acids (left) or in R5 with or without L-ascorbic acid. Cells grown in R5 without L-ascorbic acid grew significantly less than in the presence of L-ascorbic acid (n=5, q-value <0.001 at days 21 to 28 (FDR <5%).

Third row: Cells were grown in R8 containing decreasing amount of lipids (left) or in R5 with or without lipids (right) (n=2, q-value <0.001 at days 21 to through 28). In both cases grew better in the presence of lipids.

Last row: Cells were grown in R8 containing decreasing amount of Sodium selenite (left) or in R5 with or without sodium selenite. In both cases there were no major differences in cell growth.

**Figure S5: MNC RED protocol:** data from the three individuals tested in Figure 5B were averaged. The difference between the two protocols is significant at days 21 and 25 and almost significant at days 28.The graph illustrates the average fold amplification (± S.D.).

#### Tables:

| | | |
|---|---|---|
| **Culture media and supplements** | **RIT**<br>RPMI 1640<br>β-mercapto-ethanol 0.1mM<br>Ethanolamine 1.6mM<br>Lipids (1/200)<br>Insulin 10μg/mL<br>Transferrin 200μg/mL<br>(hAB-serum 10%)<br><br>***R8***<br>RPMI 1640<br>Ethanolamine 1.6mM<br>L-ascorbic acid 220 μM<br>Lipids 1/200<br>Sodium Chloride 1g/L<br>Sodium Selenite 74nM<br>Insulin 10ug/mL<br>Transferrin 200μg/mL<br><br>***R5***<br>RPMI 1640<br>L-ascorbic acid 220 μM<br>Lipids 1/200<br>Insulin 10ug/mL<br>Transferrin 200μg/mL<br><br>***R6***<br>RPMI 1640<br>L-ascorbic acid 220 μM<br>Lipids 1/200<br>Insulin 10ug/mL<br>Optiferrin 20 ug/mL<br>FeIII-EDTA 4μM<br><br>***IMIT***<br><br>IMDM with 1mM Glutamine<br>Methyl-β-Cyclodextrin 0.1mg/mL<br>Trolox50μM<br>Insulin 10μg/mL<br>Optiferrin 20 ug/mL<br>FeIII-EDTA 4μM<br>Lipids (1.5X)<br>Ethanolamine | **SED**<br>SCF 100ng/mL<br>EPO 4U/mL<br>IBMX 50 μM<br>Dexamethasone 1 μM<br><br>**SER**<br>SCF 50ng/mL<br>EPO 4U/mL<br>RU486 1 μM<br><br>**R**<br>RU486 1 μM<br><br>**Week 1 (W1)**<br>StemSpan SFEM<br>Hydrocortisone 1μM<br>SCF 50 ng/mL<br>FLT3L 16.7 ng/mL<br>IL3 6.67 ng/mL<br>EPO 1.33 U/mL<br><br>**Week 2 (W2)**<br>StemSpan SFEM<br>Hydrocortisone 1μM<br>SCF 20 ng/mL<br>IGF1 20 ng/mL<br>IL3 6.67 ng/mL<br>EPO 2 U/mL<br><br>**SEII cytokines**<br>SCF 4.4 ng/mL<br>EPO 3u/mL<br>IGF1 4.4ng/mL<br>IL3 1.5 ng/mL |
| **Table S1** | | |

| Reagent | Provider | Catalog Number |
|---|---|---|
| IMDM with 1mM Glutamine | Biochrom | FG0465 |
| RPMI 1640 with 1mM Glutamine | Gibco | 61870 |
| StemSpan SFEM | Stemcell Technologies | 09650 |
| Methyl- β -Cyclodextrin | Sigma | C4555 |
| Trolox | Sigma | 238813 |
| Insulin | Sigma | I9218 |
| Chemically defined Lipids 200X | Gibco | 11905 |
| Ethanolamine | Sigma | E0135 |
| BSA | Gibco | From Kit A1000701 |
| β-mercapto-ethanol 1000X | Gibco | 21985 |
| L-ascorbic acid | Sigma | A8960 |
| Holo-Transferrin | R&D Systems/Biotechne | 2914-HT |
| Optiferrin | FisherScience | NC9954311 |
| FeIII-EDTA | Sigma | E6760 |
| SCF | Peprotech | 300-07 |
| IGF1 | Alfa Aesar | BT-106 |
| IGF2 | Alfa Aesar | BT-107 |
| EPO | Amgen | NDC 55513-126-10 |
| Dexamethasone | Sigma | D4902 |
| RU486 | Sigma | M8046 |
| Hydrocortisone | Sigma | H0888 |
| FLT3L | Peprotech | 300-19 |
| IL3 | Peprotech | 200-03 |

**Table S2: Reagents**

A

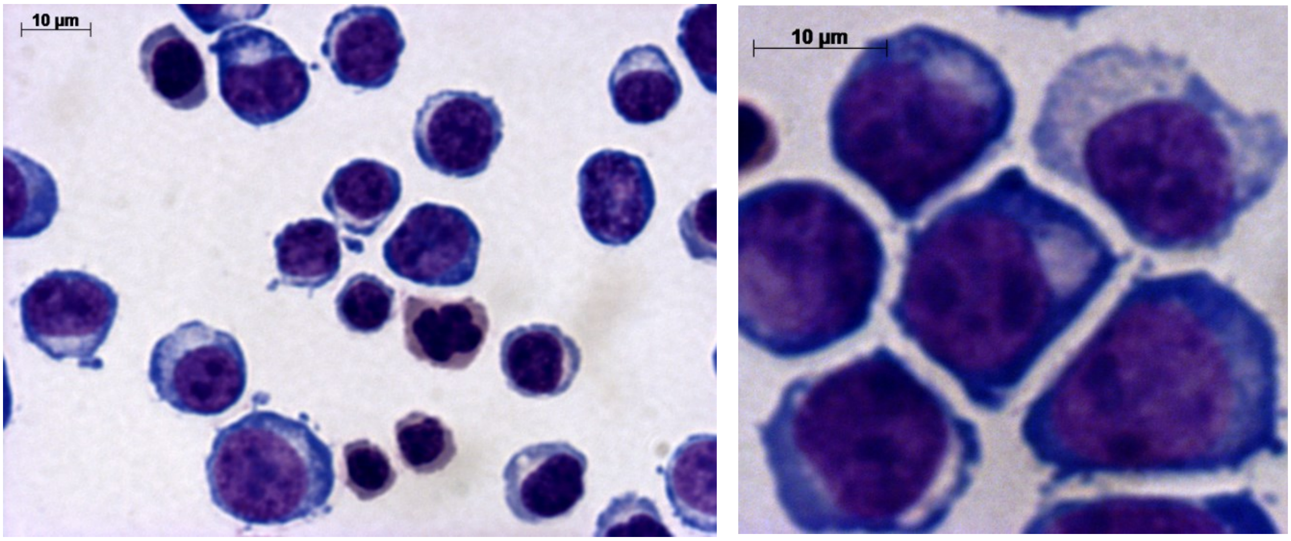
10 µm
10 µm

B

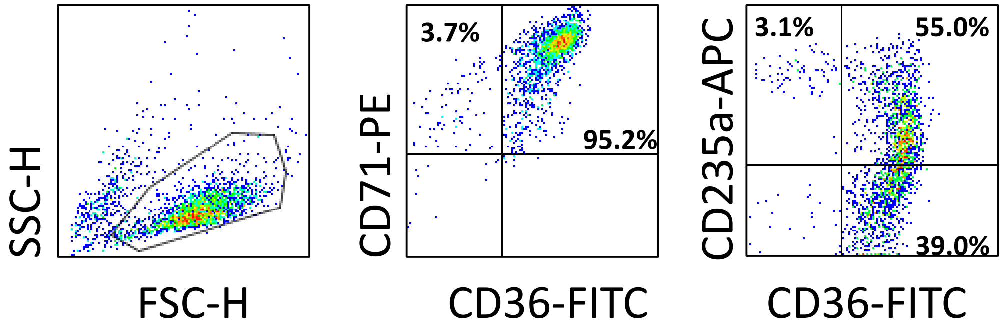
SSC-H
FSC-H
CD71-PE
3.7%
95.2%
CD36-FITC
CD235a-APC
3.1%
55.0%
39.0%
CD36-FITC

Figure S1

### p-and q-values for Figure 1A (role of dex)

| | days | p-value | q-value |
|---|---|---|---|
| 1 vs 3<br>STD vs SED | 17 | 0.0107 | 0.0168 |
| | 19 | 0.0039 | 0.0119 |
| | 21 | 0.0447 | 0.0470 |
| | 23 | 0.0153 | 0.0193 |
| | 25 | 0.0057 | 0.0119 |
| | 28 | 0.0003 | 0.0016 |
| 1 vs 4<br>STD vs SE | 17 | 0.0093 | 0.0584 |
| | 19 | 0.3093 | 0.3248 |
| | 21 | 0.0639 | 0.0806 |
| | 23 | 0.0387 | 0.0609 |
| | 25 | 0.0294 | 0.0609 |
| | 28 | 0.0319 | 0.0609 |
| 1 vs 5<br>STD vs SED | 17 | 0.0552 | 0.0232 |
| | 19 | 0.1707 | 0.0597 |
| | 21 | 0.0024 | 0.0019 |
| | 23 | 0.0000 | 0.0001 |
| | 25 | 0.0034 | 0.0019 |
| | 28 | 0.0037 | 0.0019 |
| 2 vs 3<br>(STD +SED) vs SED | 19 | 0.1219 | 0.1280 |
| | 21 | 0.0899 | 0.1180 |
| | 23 | 0.0587 | 0.1027 |
| | 25 | 0.0299 | 0.0784 |
| | 28 | 0.0005 | 0.0026 |
| 2 vs 5<br>(STD +SED )vs SER | 17 | 0.0906 | 0.0381 |
| | 19 | 0.2131 | 0.0746 |
| | 21 | 0.0036 | 0.0030 |
| | 23 | 0.0001 | 0.0003 |
| | 25 | 0.0173 | 0.0091 |
| | 28 | 0.0042 | 0.0030 |
| 3 vs 5<br>SED vs SER | 17 | 0.1289 | 0.2103 |
| | 19 | 0.2737 | 0.2874 |
| | 21 | 0.0424 | 0.1113 |
| | 23 | 0.0021 | 0.0108 |
| | 25 | 0.1602 | 0.2103 |
| | 28 | 0.3480 | 0.3045 |

Figure S2A

### p- and q-values for Figure 1C (role of dex)

| | day 17 | | day 19 | | day 21 | | day 24 | | day 28 | |
|---|---|---|---|---|---|---|---|---|---|---|
| | p-value | q-value | p-value | q-value | p-value | q-value | p-value | q-value | p-value | q-value |
| **1 vs 3** | | | | | | | | | | |
| **proE** | 0.5028 | 0.6534 | 0.1284 | 0.4026 | 0.0095 | 0.0150 | 0.1778 | 0.1400 | | |
| **basoE** | 0.6223 | 0.6534 | 0.2576 | 0.4026 | 0.0068 | 0.0150 | 0.0002 | 0.0007 | 0.1778 | 0.1467 |
| **polyE** | 0.3257 | 0.6534 | 0.3067 | 0.4026 | 0.2640 | 0.1733 | 0.0132 | 0.0208 | 0.0012 | 0.0006 |
| **orthoE** | 0.4414 | 0.6534 | 0.2877 | 0.4026 | 0.2751 | 0.1733 | 0.6035 | 0.3802 | 0.0080 | 0.0028 |
| **Retic** | 0.5415 | 0.6534 | 0.6302 | 0.6617 | 0.1133 | 0.1190 | 0.0454 | 0.0477 | 0.0011 | 0.0006 |
| **1 vs 4** | | | | | | | | | | |
| **proE** | ND | ND | ND | ND | ND | ND | | | | |
| **basoE** | ND | ND | ND | ND | ND | ND | 0.0000 | 0.0001 | | |
| **polyE** | ND | ND | ND | ND | ND | ND | 0.0027 | 0.0029 | 0.6636 | 0.2323 |
| **orthoE** | ND | ND | ND | ND | ND | ND | 0.2243 | 0.1178 | 0.0181 | 0.0095 |
| **Retic** | ND | ND | ND | ND | ND | ND | 0.0389 | 0.0272 | 0.0138 | 0.0095 |
| **1 vs 5** | | | | | | | | | | |
| **proE** | 0.4868 | 0.8520 | 0.2057 | 0.4704 | 0.0002 | 0.0006 | 0.1037 | 0.1452 | | |
| **basoE** | 0.8464 | 0.8887 | 0.2431 | 0.4704 | 0.0038 | 0.0060 | 0.0394 | 0.0828 | 0.3559 | 0.2803 |
| **polyE** | 0.4266 | 0.8520 | 0.3205 | 0.4704 | 0.2901 | 0.1828 | 0.0002 | 0.0009 | 0.0240 | 0.0378 |
| **orthoE** | 0.7996 | 0.8887 | 0.3584 | 0.4704 | 0.1342 | 0.1409 | 0.5027 | 0.4223 | 0.0519 | 0.0545 |
| **Retic** | 0.4366 | 0.8520 | 0.7888 | 0.8282 | 0.2516 | 0.1828 | 0.1498 | 0.1573 | 0.0087 | 0.0274 |
| **2 vs 3** | | | | | | | | | | |
| **proE** | 0.5065 | 0.6808 | 0.0447 | 0.1174 | | | 0.2722 | 0.2481 | | |
| **basoE** | 0.6483 | 0.6808 | 0.0098 | 0.0513 | 0.0182 | 0.0064 | 0.0002 | 0.0008 | 0.2722 | 0.1429 |
| **polyE** | 0.5203 | 0.6808 | 0.9033 | 0.9485 | 0.0028 | 0.0026 | 0.0817 | 0.1715 | 0.0097 | 0.0155 |
| **orthoE** | 0.4851 | 0.6808 | 0.1338 | 0.2341 | 0.0049 | 0.0026 | 0.1408 | 0.1971 | 0.2631 | 0.1429 |
| **Retic** | | | 0.4544 | 0.5964 | 0.4085 | 0.1072 | 0.2953 | 0.2481 | 0.0147 | 0.0155 |
| **2 vs 4** | | | | | | | | | | |
| **proE** | ND | ND | ND | ND | ND | ND | | | | |
| **basoE** | ND | ND | ND | ND | ND | ND | 0.0002 | 0.0002 | | |
| **PolyE** | ND | ND | ND | ND | ND | ND | 0.0190 | 0.0066 | 0.6876 | 0.7220 |
| **orthoE** | ND | ND | ND | ND | ND | ND | 0.0031 | 0.0016 | 0.3999 | 0.6299 |
| **Retic** | ND | ND | ND | ND | ND | ND | 0.7182 | 0.1885 | 0.0506 | 0.1593 |
| **2 vs 5** | | | | | | | | | | |
| **proE** | 0.6023 | 0.9459 | 0.0115 | 0.0181 | 0.0007 | 0.0003 | 0.1830 | 0.2402 | | |
| **basoE** | 0.9009 | 0.9459 | 0.0077 | 0.0181 | 0.0068 | 0.0018 | 0.0820 | 0.1435 | 0.4366 | 0.6112 |
| **polyE** | 0.5664 | 0.9459 | 0.8187 | 0.5362 | 0.0003 | 0.0002 | 0.0215 | 0.0565 | 0.1658 | 0.3481 |
| **orthoE** | 0.7742 | 0.9459 | 0.1463 | 0.1536 | 0.0000 | 0.0000 | 0.0103 | 0.0541 | 0.7418 | 0.7789 |
| **Retic** | | | 0.8512 | 0.5362 | 0.1109 | 0.1233 | 0.7606 | 0.7986 | 0.0230 | 0.0965 |
| **3vs 5** | | | | | | | | | | |
| **proE** | 0.8479 | 0.9013 | 0.4343 | 0.9583 | 0.0133 | 0.0700 | 0.3753 | 0.5254 | | |
| **basoE** | 0.5874 | 0.9013 | 0.9126 | 0.9583 | 0.2320 | 0.3046 | 0.9999 | 0.8400 | 0.3552 | 0.1865 |
| **polyE** | 0.8584 | 0.9013 | 0.8093 | 0.9583 | 0.0878 | 0.1536 | 0.8922 | 0.8400 | 0.0571 | 0.0400 |
| **orthoE** | 0.7843 | 0.9013 | 0.7706 | 0.9583 | 0.7087 | 0.7441 | 0.1679 | 0.3527 | 0.0136 | 0.0143 |
| **retic** | | | 0.5149 | 0.9583 | 0.0796 | 0.1536 | 0.0059 | 0.0246 | 0.0038 | 0.0080 |

Figure S2B

### p- and q-values for Figures 2 and S4 (R5 and R8 )

| | days | p-value | q-value |
|---|---|---|---|
| Ethanolamine in R8 1.6mM vs 0mM | 17 | 0.521698 | 0.273891 |
| | 21 | 0.000043 | 0.000045 |
| | 23 | 0.000546 | 0.000382 |
| | 25 | 0.000004 | 0.000008 |
| R5 vs R5 - acid ascorbic | 16 | 0.000000 | 0.000000 |
| | 19 | 0.006416 | 0.004491 |
| | 21 | 0.014597 | 0.007663 |
| | 23 | 0.029402 | 0.012349 |
| | 26 | 0.050052 | 0.017518 |
| | 28 | 0.004295 | 0.004491 |
| R5 vs R5 - lipids | 17 | 0.516746 | 0.180861 |
| | 19 | 0.006330 | 0.002658 |
| | 21 | 0.000077 | 0.000040 |
| | 23 | 0.000001 | 0.000001 |
| | 25 | 0.000004 | 0.000004 |
| | 28 | 0.000035 | 0.000025 |
| R5 versus R8 | 16 | 0.81292 | 0.24388 |
| | 19 | 0.00003 | 0.00006 |
| | 21 | 0.00052 | 0.00055 |
| | 23 | 0.00147 | 0.00103 |
| | 26 | 0.00375 | 0.00158 |
| | 28 | 0.00342 | 0.00158 |

Figure S2C

### p- and q-value for Figure 3A (iron chelator)

| | days | p-value | q-value |
|---|---|---|---|
| 1 vs 2<br>200 vs 50 μg/mL holo-Tf | 16 | 0.8053 | 0.1691 |
| | 19 | 0.0048 | 0.0012 |
| | 21 | 0.0000 | 0.0000 |
| | 23 | 0.0000 | 0.0000 |
| | 28 | 0.0000 | 0.0000 |
| 1 vs 3<br>200 μg/mL holo-Tf<br>vs<br>50 μg/mL holo-Tf +3 μM ATA | 16 | 0.9437 | 0.5945 |
| | 19 | 0.3285 | 0.2587 |
| | 21 | 0.0010 | 0.0033 |
| | 23 | 0.0051 | 0.0080 |
| | 28 | 0.0563 | 0.0591 |
| 2 vs 3<br>50 μg/mL holo-Tf<br>vs<br>50 μg/mL holo-Tf +3 μM ATA | 16 | 0.7752 | 0.1628 |
| | 19 | 0.0012 | 0.0003 |
| | 21 | 0.0000 | 0.0000 |
| | 23 | 0.0000 | 0.0000 |
| | 28 | 0.0000 | 0.0000 |
| 2 vs 4<br>50 μg/mL holo-Tf<br>vs<br>50 μg/mL holo-Tf +3 μM Eryochrome cyanide | 16 | 0.4411 | 0.0926 |
| | 19 | 0.0007 | 0.0002 |
| | 21 | 0.0000 | 0.0000 |
| | 23 | 0.0000 | 0.0000 |
| | 28 | 0.0000 | 0.0000 |
| 2 vs 5<br>50 μg/mL holo-Tf<br>vs<br>50 μg/mL holo-Tf +3 μM FeIII-EDTA | 16 | 0.8456 | 0.3551 |
| | 19 | 0.0651 | 0.0342 |
| | 21 | 0.0009 | 0.0007 |
| | 23 | 0.0001 | 0.0001 |
| | 28 | 0.0011 | 0.0007 |

Figure S2D

### p- and q- values for Figure 3B (Optiferrin titration)

| | days | p-value | q-value |
|---|---|---|---|
| Optiferrin 50 mg/mL vs 6.25 mg/mL | 19 | 0.1441 | 0.1513 |
| | 21 | 0.0970 | 0.1357 |
| | 24 | 0.0862 | 0.1357 |
| | 28 | 0.2715 | 0.2280 |
| | 31 | 0.0034 | 0.0142 |
| Optiferrin 50mg/mL vs 0 mg/mL | 19 | 0.0454 | 0.0477 |
| | 21 | 0.0215 | 0.0282 |
| | 24 | 0.0057 | 0.0123 |
| | 28 | 0.0070 | 0.0123 |
| | 31 | 0.0010 | 0.0053 |
| Optiferrin 25mg/mL VS 0 mg/mL | 19 | 0.0823 | 0.0173 |
| | 21 | 0.0261 | 0.0069 |
| | 24 | 0.0066 | 0.0023 |
| | 28 | 0.0045 | 0.0023 |
| | 31 | 0.0037 | 0.0023 |
| Optiferrin 12.5mg/mL vs 0 mg/mL | 19 | 0.1043 | 0.0219 |
| | 21 | 0.0379 | 0.0100 |
| | 24 | 0.0091 | 0.0048 |
| | 28 | 0.0073 | 0.0048 |
| | 31 | 0.0146 | 0.0051 |
| Optiferrin 6.5mg/mL vs 0 mg/mL | 19 | 0.1259 | 0.0529 |
| | 21 | 0.0532 | 0.0280 |
| | 24 | 0.0159 | 0.0111 |
| | 28 | 0.0126 | 0.0111 |
| | 31 | 0.0019 | 0.0041 |
| 0mg/mL vs 50mg/mL (Holo-Tf) | 19 | 0.0681 | 0.0179 |
| | 21 | 0.0124 | 0.0061 |
| | 24 | 0.0048 | 0.0051 |
| | 28 | 0.0174 | 0.0061 |

Figure S2E

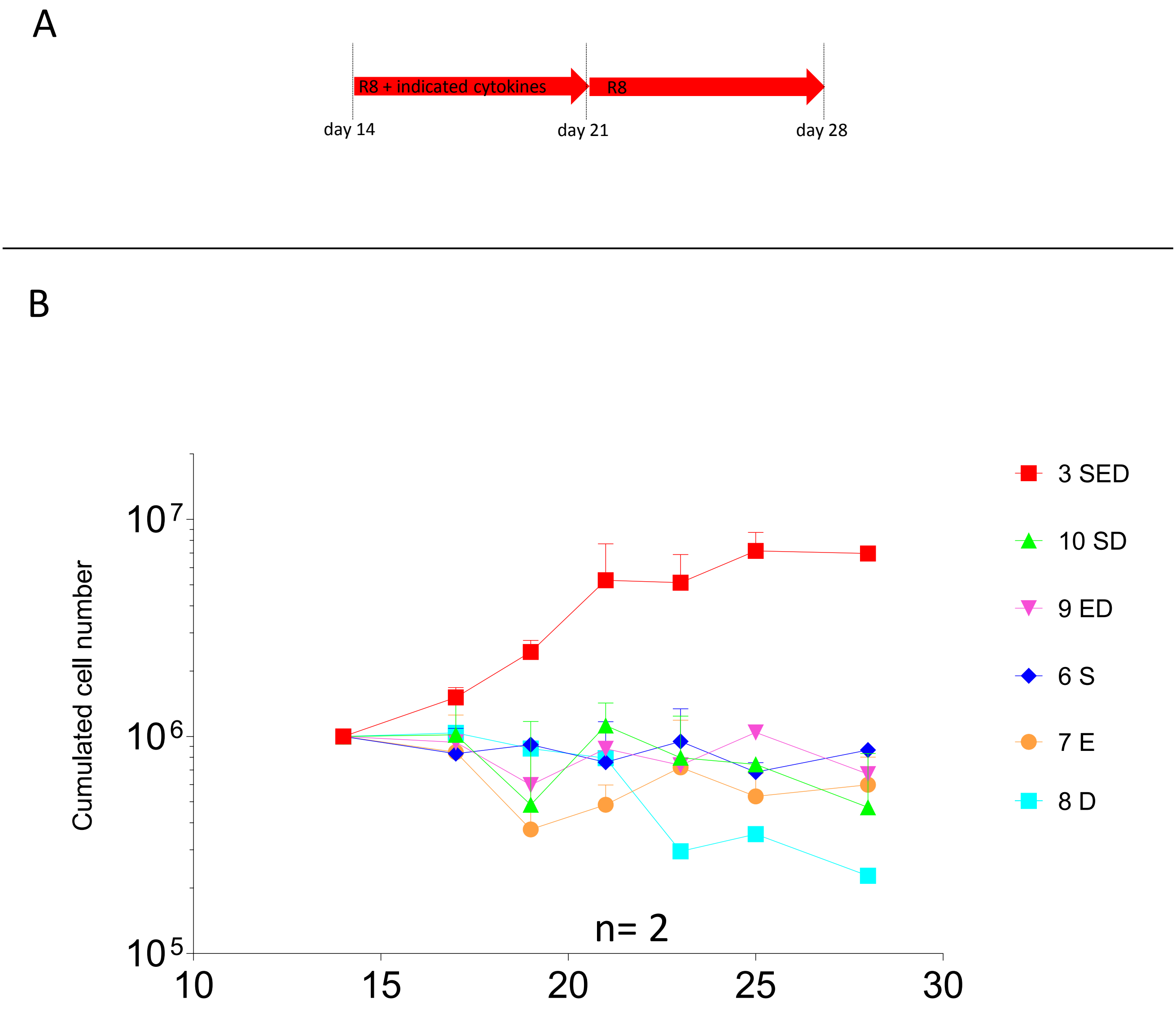

A
R8 + indicated cytokines
R8
day 14
day 21
day 28
B
Cumulated cell number
$10^7$
$10^6$
$10^5$
10
15
20
25
30
n= 2
3 SED
10 SD
9 ED
6 S
7 E
8 D


Figure S3

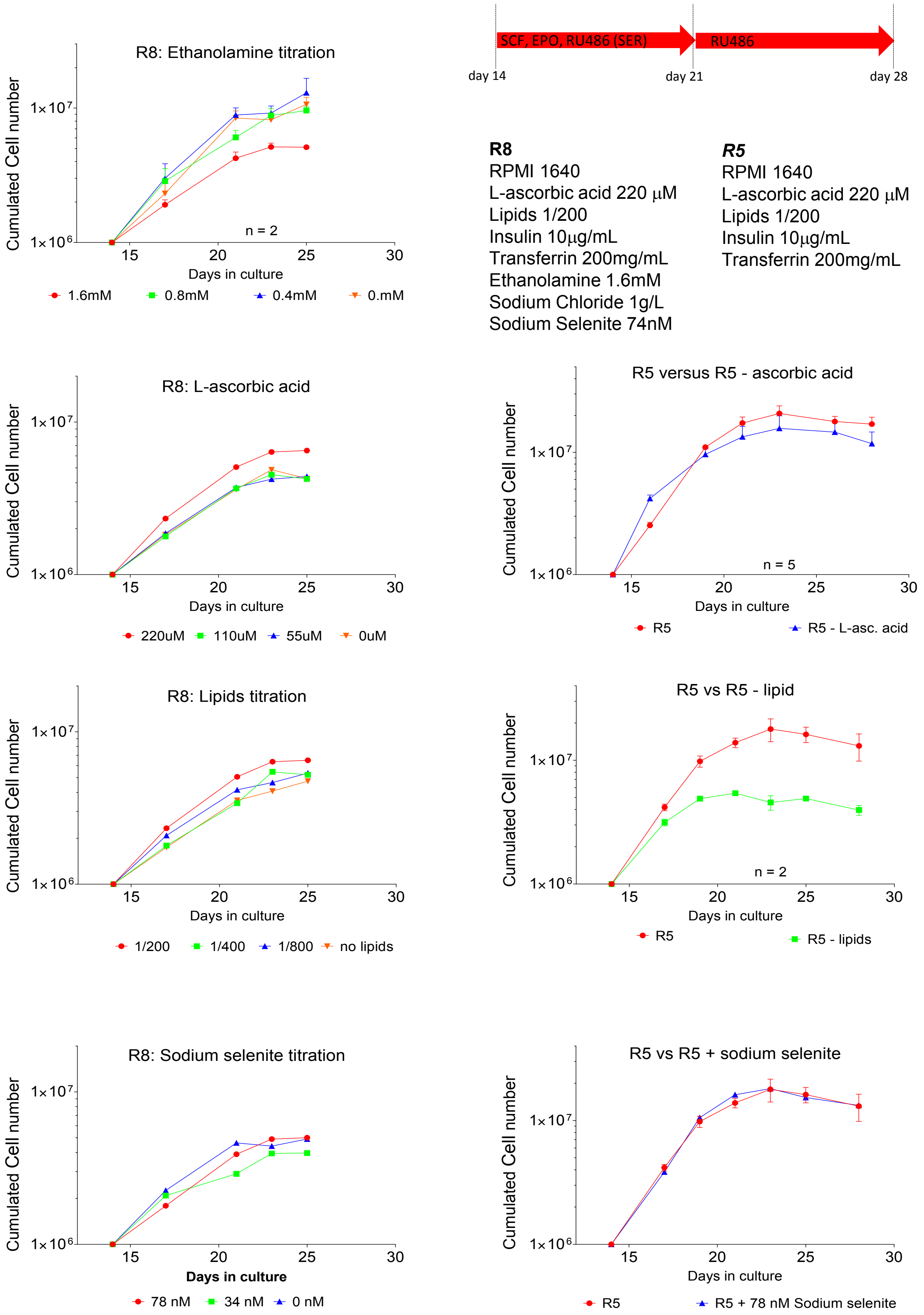
SCF, EPO, RU486 (SER)
RU486
day 14
day 21
day 28
R8: Ethanolamine titration
Cumulated Cell number
1×10^7
1×10^6
n = 2
15
20
25
30
Days in culture
1.6mM
0.8mM
0.4mM
0.mM
R8
RPMI 1640
L-ascorbic acid 220 μM
Lipids 1/200
Insulin 10μg/mL
Transferrin 200mg/mL
Ethanolamine 1.6mM
Sodium Chloride 1g/L
Sodium Selenite 74nM
R5
RPMI 1640
L-ascorbic acid 220 μM
Lipids 1/200
Insulin 10μg/mL
Transferrin 200mg/mL
R8: L-ascorbic acid
220uM
110uM
55uM
0uM
R5 versus R5 - ascorbic acid
n = 5
R5
R5 - L-asc. acid
R8: Lipids titration
1/200
1/400
1/800
no lipids
R5 vs R5 - lipid
n = 2
R5
R5 - lipids
R8: Sodium selenite titration
78 nM
34 nM
0 nM
R5 vs R5 + sodium selenite
R5
R5 + 78 nM Sodium selenite

Figure S4

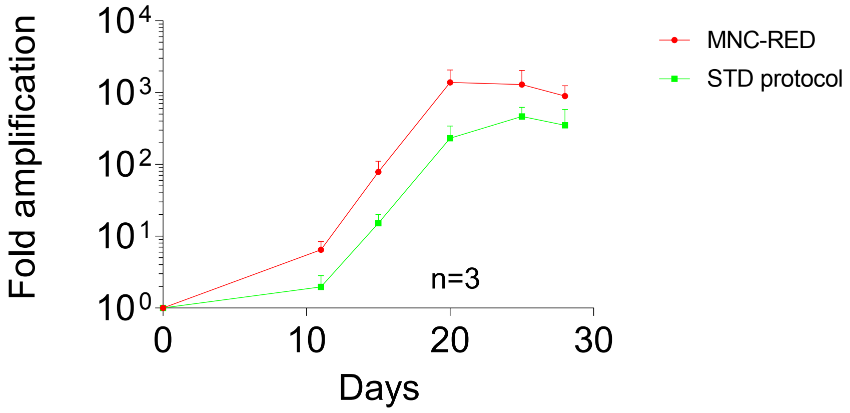


| Days | p-value | q-value |
|---|---|---|
| **15** | 0.8096 | 0.7000 |
| **20** | 0.0002 | 0.0008 |
| **25** | 0.0041 | 0.0087 |
| **28** | 0.0474 | 0.0664 |

Figure S5
